# Cross-site harmonization of diffusion MRI data without matched training subjects

**DOI:** 10.1101/2024.05.01.591994

**Authors:** Alberto De Luca, Tine Swartenbroekx, Harro Seelaar, John van Swieten, Suheyla Cetin Karayumak, Yogesh Rathi, Ofer Pasternak, Lize Jiskoot, Alexander Leemans

## Abstract

**Purpose:** Diffusion MRI (dMRI) data typically suffer of significant cross-site variability, which prevents naively performing pooled analyses. To attenuate cross-site variability, harmonization methods such as the rotational invariant spherical harmonics (RISH) have been introduced to harmonize the dMRI data at the signal level. A common requirement of the RISH method, is the availability of healthy individuals who are matched at the group level, which may not always be readily available, particularly retrospectively. In this work, we propose a framework to harmonize dMRI without matched training groups.

**Methods:** Our framework learns harmonization features while controlling for potential covariates using a voxel-based generalized linear model (RISH-GLM). RISH-GLM allows to simultaneously harmonize data from any number of sites while also accounting for covariates of interest, thus not requiring matched training subjects. Additionally, RISH-GLM can harmonize data from multiple sites in a single step, whereas RISH is performed for each site independently.

**Results:** We considered data of training subjects from retrospective cohorts acquired with 3 different scanners and performed 3 harmonization experiments of increasing complexity. First, we demonstrate that RISH-GLM is equivalent to conventional RISH when trained with data of matched training subjects. Secondly, we demonstrate that RISH-GLM can effectively learn harmonization with two groups of highly unmatched subjects. Thirdly, we evaluate the ability of RISH-GLM to simultaneously harmonize data from 3 different sites.

**Discussion:** RISH-GLM can learn cross-site harmonization both from matched and unmatched groups of training subjects, and can effectively be used to harmonize data of multiple sites in one single step.

## Introduction

Diffusion MRI (dMRI) has become a pivotal technique to investigate brain structure non-invasively^1^. The signal measured with dMRI originates from the motion of water molecules at the microscopic scale^2^. In combination with appropriate modelling^3^, dMRI allows to infer microstructural properties of tissues. For example, diffusion tensor imaging^4,5^ (DTI), one of the most commonly applied dMRI techniques, provides metrics such as the mean diffusivity (MD) and the fractional anisotropy^4,6^ (FA), which are related to the root mean square displacement and the anisotropy of the diffusion process, and reflect properties of the biological environment. Over time, these metrics have become established in the study of white matter microstructure, and found application in the study of neurological diseases^7^, including Alzheimer’s disease^8^, small vessel disease^9,10^ and frontotemporal dementia^11^, among others.

A possible limitation to the use of dMRI in clinical research is that its measurements strongly depend on the employed MRI hardware and software^12,13^. As such, metrics derived from dMRI can typically be compared only within a single site, even when acquisition parameters are kept constant. In the latest years, several methods have been proposed to tackle such cross-site variability and harmonize dMRI. Broadly speaking, the goal of harmonization methods can be summarized as removing cross-site differences while not altering the sensitivity of dMRI to biological effects of interest. Two main families of post-processing dMRI harmonization methods have been proposed to date. The first family aims to remove batch effects during the analysis steps. Methods such as ComBat^14–16^, for example, aim to remove cross-site differences on the final metrics derived from dMRI. As such, harmonization is applied independently to each considered dMRI metric, such as FA or MD. The second family of dMRI harmonization methods aims to remove cross-site effects on the acquired data before any quantification step. Notable examples hereof are the rotational invariant spherical harmonics^17,18^ (RISH) framework, or other recently proposed deep learning alternatives^19,20^. Differently from methods such as ComBat, which harmonize end-point diffusion metrics independently from each other, RISH and comparable frameworks harmonize the source dMRI data. As such, the source data is harmonized once as part of the preprocessing pipeline, and can be then reused for multiple analyses with any method of choice. For example, we have previously demonstrated that dMRI data of patients with small vessel disease^21,22^ can be effectively harmonized with RISH and then be used for both voxel-wise based analyses and connectomics.

One of the challenges when applying dMRI harmonization methods such as RISH, is that they require data of healthy participants for calibration between sites. For example, previous work^17^ has demonstrated that the RISH method requires 12 or more training subjects matched at the group level for each site to be harmonized, or repeated scans of so-called travelling heads. This can be particularly challenging to achieve for studies focusing on patient populations, where healthy controls may not always be recruited, or when retrospectively harmonizing multiple different cohorts with differences in average age or sex distribution. Furthermore, the concept of matched groups is ill-defined and study dependent. In practice, it is very often interpreted as a synonym of age and sex matching, but other factors such as education, presence of genetic mutations, among others, may affect the dMRI signal and might need to be considered for specific studies.

In this study, we propose a generalization of the RISH framework that determines cross-site harmonization while also considering potential confounders. This new approach accounts for potential biases when learning harmonization from unmatched groups of training subjects, and also allows to harmonize data from multiple sites in one single step. We demonstrate the potential of this framework through three experiments of growing complexity employing dMRI data of subjects from three different sites, while controlling for sex and age.

## Methods

### Theory: RISH and RISH-GLM

The conventional RISH harmonization method is based on the representation of diffusion MRI data acquired at a certain b-value (i.e., a diffusion “shell”) with spherical harmonics. Spherical harmonics are a set of functions on the unit sphere characterized by an order and a degree. For each order *i*, rotational invariant representations (i.e., RISH coefficients) can be determined, and are indicated as *L_i_*. Mirzaalian et al. and subsequent works^17,23^ have shown that it is possible to harmonize dMRI data by learning the scaling factors between corresponding RISH coefficients determined in two groups of matched training subjects. The same framework can be also used to harmonize multi-shell data effectively by harmonizing each shell independently^18^. The harmonization essentially determines a voxel-wise scaling factor associated with each RISH feature to map a target site to a reference site. Mathematically, the scaling coefficient *ϑ* associated to the spherical harmonics of order *i* can be written as

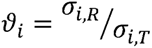

where *σ* indicate the average RISH features of the reference (R) and target sites (T). Subsequently, dMRI datasets from the target site can be harmonized as follows: 1) dMRI data is converted to spherical harmonics; 2) their RISH features calculated and multiplied by the scaling factor *ϑ_i_*; 3) harmonized dMRI data is reconstructed from the corrected spherical harmonics representation. The assumption behind this approach is that cross-site differences in *σ* purely originate from scanner-related properties. For this assumption to hold, the groups from which the *σ* values are computed must be matched across all factors that may affect the dMRI signal. In practice, matching for all relevant factors can be challenging. When this is not the case, residual systematic group differences will be assimilated into the scaling factors, potentially biasing the factors and making them less generalizable to harmonize other subjects from the same site.

To alleviate the matching requirement, we propose a new formulation based on a general linear model (GLM) that decomposes the effect of confounders such as age and sex on the RISH features:

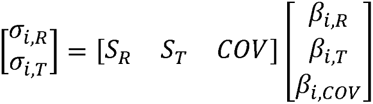

where *S* are dummy variables assuming a value of 1 when a subject belongs to a certain site, and 0 otherwise. This approach, dubbed RISH-GLM, is based on two assumptions: 1) that the relationship between RISH features and covariates is linear in the considered range; 2) limited collinearity and interaction between different covariates. Of note, linearity in the relation between covariates and RISH-features, does not imply nor require linearity between covariates and diffusion metrics (e.g., fractional anisotropy). The RISH-GLM formulation allows to alleviate the need to match the average demographics at the group level across all sites. In practice, however, a reasonably even distribution of the effects of interest across sites is still needed to reliably estimate the associated coefficients *β*, from which the scaling coefficients can be determined as 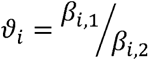. The formulation above can readily be extended to account for multiple sites by considering additional columns on the right side as follows:

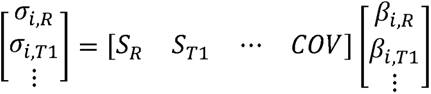

### MRI data

We retrospectively collected data from the Frontotemporal Dementia Risk Cohort^24^ (FTD-RISC), which is a longitudinal study following up individuals at risk of frontotemporal dementia (FTD) curated at the Erasmus MC University Medical Center in Rotterdam (the Netherlands). In FTD-RISC, three major combinations of MRI hardware and software have been employed, which we refer to as Site 1, Site 2, Site 3. Data from all sites are used throughout this study to demonstrate the potential of GLM-RISH. T1- weighted MRI data were acquired at all sites with a resolution of 1×1×1mm^3^ isotropic. dMRI data were acquired at all sites with a similar dMRI protocol featuring 1 b = 0 s/mm^2^ volume, and 60 gradient directions at b = 1000 s/mm^2^. Data of Site 1 were acquired with a 3T Philips Achieva scanner located at Leiden University Medical Center (the Netherlands) running on software release R5 with a 32 channels head coil. At this site, key imaging parameters were voxel size 2×2×2mm^3^, TE=80ms, TR=9.25ms. Data of Site 2 were acquired with a GE Healthcare MR750 3T scanner located at Erasmus Medical Center (the Netherlands) with a 48 channels head coil. At this site, key imaging parameters were voxel size 2×2×2mm^3^, TE=57ms, TR=9s. Data of Site 3 were acquired with an older version of the scanner at Site 1, running on software release R4 and equipped with an 8 channels head coil. At this site, dMRI data were acquired with voxel size 2×2×2mm^3^, TE=80ms, TR=8.25s. For this study, only data of healthy control subjects (non-carriers) and pathogenic variant carriers who did not have FTD-related symptoms at the time of scanning were considered.

### MRI processing

MRI data was processed with a previously presented automated processing pipeline^25^. T1-weighted data was processed with CAT12 (https://neuro-jena.github.io/cat/) to remove bias fields and perform skull stripping. dMRI data was processed with MRIToolkit (https://github.com/delucaal/MRIToolkit) and ExploreDTI^26,27^ to perform signal drift correction^28^, then denoised with the MPPCA method^29^. Subsequently, corrections for Gibbs ringing^30^, motion and EPI corrections were performed, including b-matrix rotation^31^ EPI distortions were corrected by means of a non-linear registration to the T1-weighted data at a 2mm resolution. Robust estimation of the diffusion tensor was performed using REKINDLE^32^, then the fractional anisotropy (FA) and the mean diffusivity (MD) were calculated^5^. Quality assessment of the data was performed by generating summary screenshots of FA, MD and fit residuals of each dataset, which were visually inspected by a trained researcher (ADL). Data exhibiting excessive motion or apparent artifacts (e.g., due to braces or other interference sources) were discarded.

### Harmonization experiments

We refer to RISH[matched/unmatched] and RISH-GLM[matched/unmatched] to indicate whether the methods were trained with matched or unmatched groups of participants from different sites, respectively. To perform RISH and RISH-GLM harmonization, spherical harmonics were fitted to dMRI data using in-house software written in MATLAB, which implemented L2 regularized least squares. RISH features were then calculated and spatially normalized to a study specific template as previously proposed ^17^. The code used to perform the aforementioned steps is openly available at https://github.com/delucaal/RISH-GLM.

We evaluated the performance of RISH-GLM as compared to conventional RISH with three experiments of growing complexity. Table 1 shows an overview of the datasets used in the following three experiments.

**Table 1:**
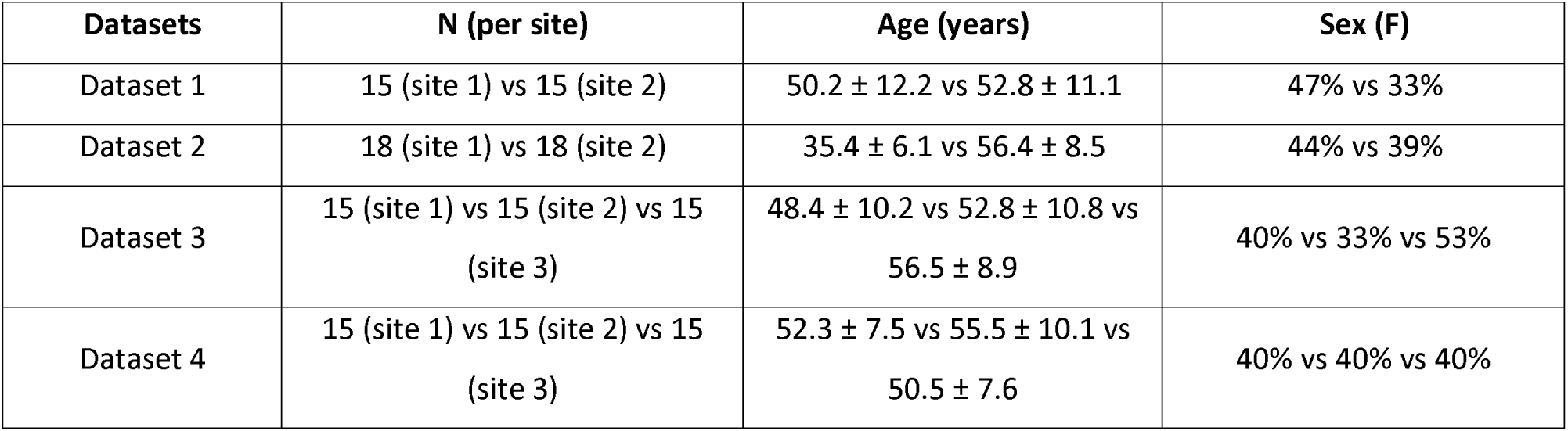
An overview of the datasets used to train RISH and RISH-GLM in the three experiments. N = sample size of each site. F = female.

| Datasets | N (per site) | Age (years) | Sex (F) |
| --- | --- | --- | --- |
| Dataset 1 | 15 (site 1) vs 15 (site 2) | $50.2 \pm 12.2$ vs $52.8 \pm 11.1$ | 47% vs 33% |
| Dataset 2 | 18 (site 1) vs 18 (site 2) | $35.4 \pm 6.1$ vs $56.4 \pm 8.5$ | 44% vs 39% |
| Dataset 3 | 15 (site 1) vs 15 (site 2) vs 15 (site 3) | $48.4 \pm 10.2$ vs $52.8 \pm 10.8$ vs $56.5 \pm 8.9$ | 40% vs 33% vs 53% |
| Dataset 4 | 15 (site 1) vs 15 (site 2) vs 15 (site 3) | $52.3 \pm 7.5$ vs $55.5 \pm 10.1$ vs $50.5 \pm 7.6$ | 40% vs 40% vs 40% |

In experiment 1, we investigate whether RISH-GLM is equivalent to the conventional RISH method when trained on data of 15 individuals matched at the group level for both age and sex from Dataset 1. For each spherical harmonics’ order, the scaling factors were computed with both RISH and RISH-GLM accounting for both age and sex, and visually compared. Subsequently, the training data from Site 2 was harmonized, and the diffusion tensor model was fit ^32^as mentioned in the previous section. FA maps were computed and registered to a common space using the TBSS_PNL pipeline (https://github.com/pnlbwh/tbss). In short, this pipeline is based on the FSL TBSS^33^ approach, but replaces the registration steps with ANTS^34^ for additional spatial accuracy. Boxplots of FA computed on the white matter skeleton were compared before and after harmonization. Given that FA is known to be strongly associated with age^35^, we additionally visualized their relation by means of scatterplots, and evaluated their Pearson correlation coefficient before and after harmonization. As no ground truth exists for age and sex effects, we evaluated the agreement between the effects estimated by RISH-GLM on the whole datasets and age and sex effects within the individual sites by means of Pearson correlations. To estimate the effects within the individual sites, general linear models accounting for age, sex and intercept were estimated at the voxel-level for site 1 and site 2 independently.

Subsequently, in experiment 2, we evaluated the feasibility of removing scanner-related cross-site differences by learning harmonization with two age-unmatched groups of subjects from Dataset 2. RISH- GLM was trained by providing age and sex of each individual as covariates. As in the previous experiment, we compared the scaling factors corresponding to different RISH features, and visually inspected the spatial maps corresponding to age and sex effects. Harmonization was applied to the training set (Dataset 2). Boxplots of FA and scatterplots of FA as a function of age were derived as explained in experiment 1. Considering the large age difference between the two groups in Dataset 2, we anticipated to observe differences in FA between the two groups before harmonization. We expected such differences not to be completely removed after harmonization if the applied method truly removes only scanner-related differences and not biological effects. Furthermore, we expected a successful harmonization to recover a Pearson correlation coefficient between age and FA as similar as possible to the one determined in experiment 1. The same analyses were repeated on an independent dataset (Dataset 1) for validation purposes. We anticipated that RISH-GLM harmonization trained with age-unmatched subjects (Dataset 2) would generalize to age-matched subjects (Dataset 1), whereas RISH harmonization would not.

In experiment 3, we evaluated whether the RISH-GLM framework allows to effectively harmonize data from multiple sites from Dataset 3 in a single step. FA maps were calculated and aligned to a common space as previously explained. Next to boxplots of FA derived on the white matter skeleton, we additionally performed voxel-wise t-tests while correcting for age and sex to also evaluate the effectiveness of RISH-GLM at removing potential regional cross-site differences. Comparisons between the three sites were performed pairwise (i.e., Site1 vs Site 2 and Site 3, Site 2 vs Site 3) using FSL randomise with threshold free cluster enhancement. Finally, to validate the one-step RISH-GLM harmonization on independent subjects, we applied the trained harmonization to age and sex-matched subjects of dataset 4, which shared only 2 out of 45 subjects with dataset 3. Here we evaluated the effectiveness of RISH-GLM at removing cross-site differences by means of boxplots of FA calculated on the white matter skeleton, and by evaluating the relation between FA and age as in the previous experiments.

In all experiments, Site1 was used as reference and its values should remain constant. However, boxplots of the reference site might exhibit slight differences because of how the TBSS_PNL pipeline – used to compute skeletonized values for the boxplots – inherently works. TBSS_PNL constructs a white matter skeleton based on all input data (both reference and target sites). Since harmonization alters the target site’s data, the global skeleton changes slightly, which can introduce minor shifts in the reference site’s FA distributions

## Results

### Experiment 1: RISH-GLM and RISH with matched training subjects (expect no differences)

Figure 1 shows the scaling maps *ϑ_i_* corresponding to spherical harmonics up to order 6 that were calculated with RISH and RISH-GLM using training subjects matched for age and sex at the group level from two different sites (hence the “[Matched]” suffix). The maps calculated by both methods are remarkably similar, as shown by the corresponding relative difference maps. The average relative differences between RISH[Matched] and RISH-GLM[Matched] are 0.38% for *ϑ*_0_, 0.73% for *ϑ*_2_, 1.32% for *ϑ*_4_, and 1.45% for *ϑ*_6_. While differences are globally less than 2%, local differences up to 20% can be appreciated in the bottom row of Figure 1. Such differences are mostly located in areas of partial volume with cerebrospinal fluid or grey matter, but not in deep white not in white matter, and might originate from small imbalances in covariates or site-specific artefacts. Indeed, age and sex maps estimated with RISH-GLM (Supporting Information Fig. S1) show that these two covariates can have a small but non-zero spatially varying effect on RISH scales, even when these are calculated in two matched groups.

**Figure 1:**
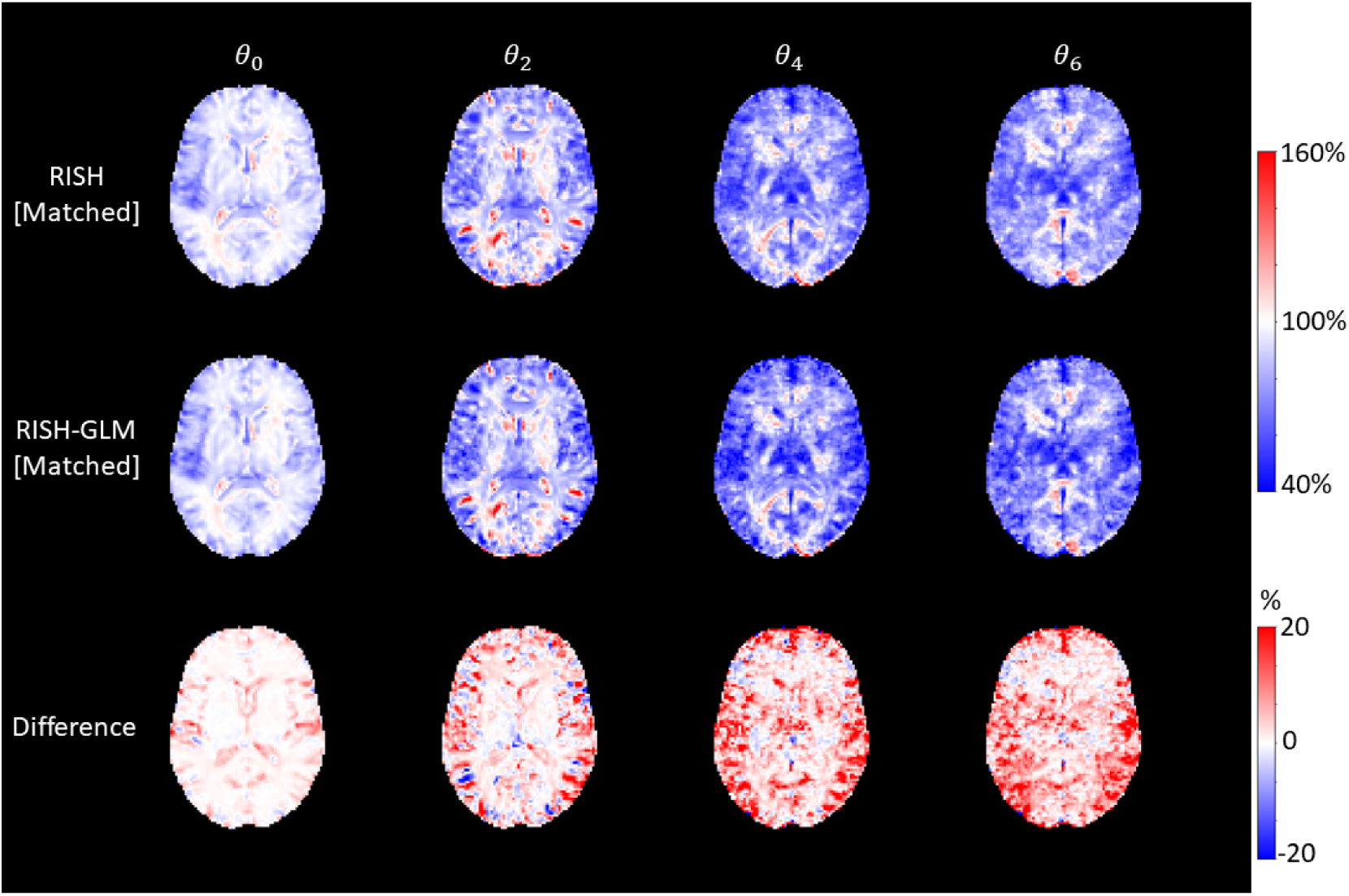
Voxel-wise scaling maps calculated in dataset 1 using group level-matched training subjects with RISH (first row), RISH-GLM (second row) and their difference (last row) for different orders of spherical harmonics (columns). Scaling maps calculated from both methods are globally similar, but local differences up to ±20% can be noticed particularly at the interface between grey matter and cerebrospinal fluid, and in the deep white matter for *θ*_4_ and *θ*_6_.

In dataset 1, age and sex effects estimated with RISH-GLM in white matter are generally in agreement with age effects calculated within site 1 and site 2 independently. The correlation coefficient between age effects estimated in the whole cohort and in sites 1/2 are 0.80/0.74 for *ϑ*_0_, 0.86/0.87 for *ϑ*_2_, 0.62/0.83 for *ϑ*_4_, 0.55/0.83 for *ϑ*_6_, respectively. The correlation coefficient for sex effects estimated in the whole cohort and in sites 1/2 are 0.75/0.70 for *ϑ*_0_, 0.72/0.70 for *ϑ*_2_, 0.53/0.89 for *ϑ*_4_, 0.56/0.89 for *ϑ*_6_, respectively.

Figure 2 shows boxplots of average FA values of Site 1 and Site 2 obtained on the white matter skeleton before harmonization, and after harmonization with RISH and RISH-GLM. Before harmonization, a relative difference in average FA values between Site 1 and Site 2 equal to −9.7% (significant t-test, p≤0.05) can be observed, which is effectively removed by both RISH (0.5%, p=0.80) and RISH-GLM (1.7%, p=0.37). The second row of Figure 2 shows the existence of a negative relation between age and FA within individual sites. When pooling unharmonized data together (first row), a biased correlation coefficient (rho) equal to −0.5 can be observed. After harmonization with both RISH and RISH-GLM, a linear negative relation between age and FA can be observed with equal correlation coefficients rho = −0.78. In both cases, the measurements from each site are well distributed around the regression line.

**Figure 2:**
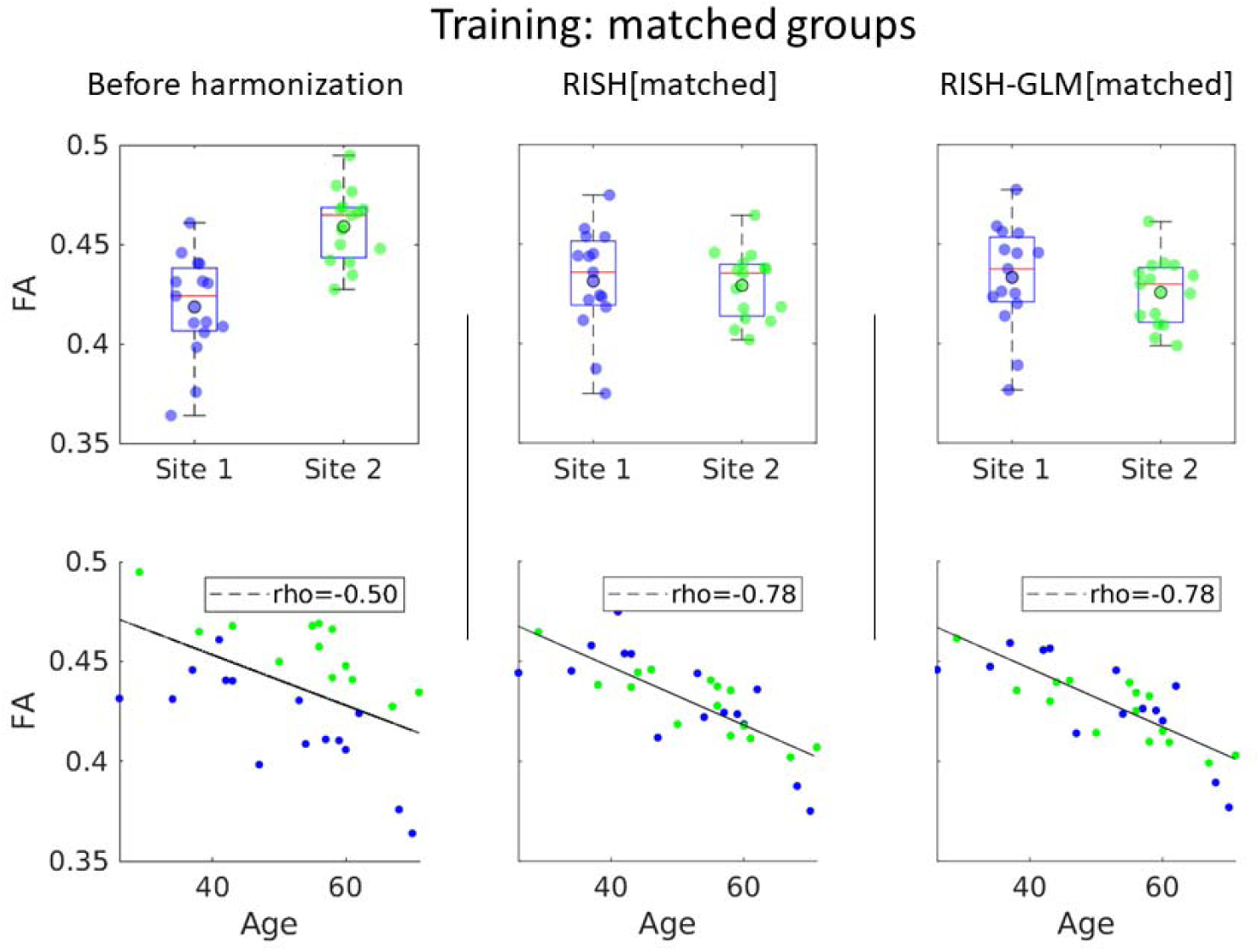
The top row shows boxplots of average FA values in the white matter skeleton per site of dataset 1, before harmonization (first column), after harmonization with RISH (middle column) and with RISH-GLM (last column). The second row shows scatterplots of the same average FA values as a function of age. Harmonization with both RISH and RISH-GLM was trained with group level-matched subjects.

### Experiment 2: RISH-GLM and RISH with unmatched training subjects

Figure 3 shows the scaling maps calculated for different orders of spherical harmonics with RISH and RISH-GLM trained with two groups of subjects that were not matched at the group level (hence the suffix “[Unmatched]”). In contrast to Figure 1, where RISH and RISH-GLM produced similar scaling maps, large differences between the two methods can be observed when using unmatched training subjects, particularly for higher orders of spherical harmonics. Compared to the reference scaling maps calculated with RISH with matched subjects (Figure 1), scaling maps calculated with unmatched subjects with RISH show large differences particularly around the ventricles, and at the interface between grey matter and cerebrospinal fluid for *ϑ*_0_ and *ϑ*_2_. For higher spherical harmonics orders (i.e., *ϑ*_4_, *ϑ*_6_), widespread differences in both the white and the grey matter can be observed. Compared to the reference (RISH[matched]), average relative differences of RISH[unmatched] and RISH-GLM[unmatched] are −10.2% and −3.3% for *ϑ*_0_, −12.0% and −2.6% for *ϑ*_2_, −10.5% and 2.9% for *ϑ*_4_, −10.6% and 3.1% for *ϑ*_6_, respectively, highlighting that scaling maps computed with RISH-GLM[unmatched] are on average closer to the reference(RISH[matched]) than RISH[unmatched].

**Figure 3:**
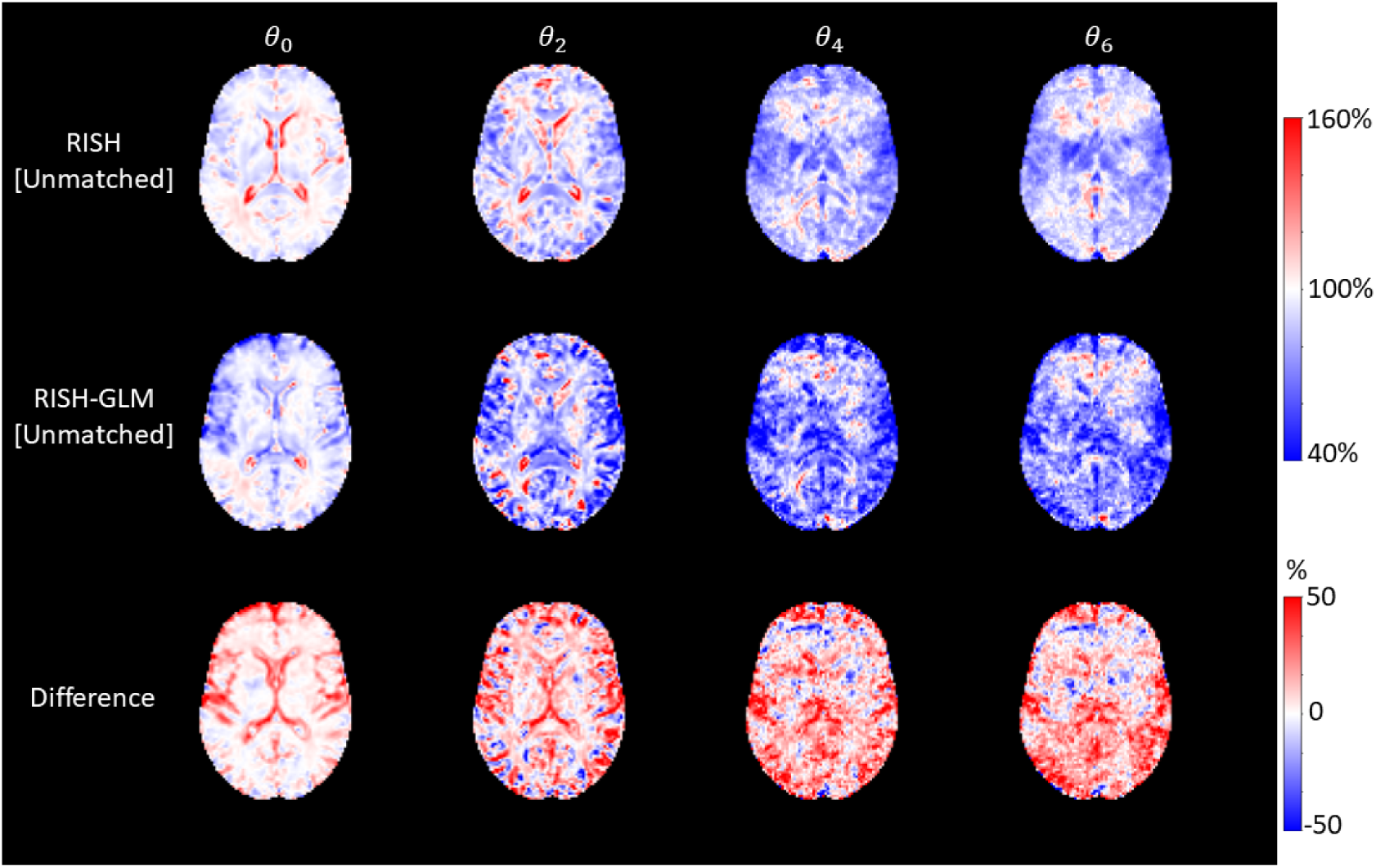
Voxel-wise scaling maps calculated using group age-unmatched training subjects (dataset 2) with RISH (first row), RISH-GLM (second row) and their difference (last row) for different orders of spherical harmonics (columns). Compared to Figure 1 (matched training groups), scaling maps calculated show different patterns with local differences up to ±50%.

Figure 4 shows the spatial coefficients of age and sex effects associated to different spherical harmonics orders. Considering *ϑ*_0_, it can be appreciated that age has a large effect around the ventricles and at the interface between grey matter and cerebrospinal fluid. At higher orders, age has a stronger effect on the brain white matter, particularly in the corpus callosum and corticospinal tracts. Similar observations also hold for sex effects. Importantly, both covariates have the largest effects in areas where RISH[matched] and RISH-GLM[unmatched] differ the most. In dataset 2, age and sex effects estimated with RISH-GLM in white matter are generally in agreement with age effects calculated within groups of Dataset 2 independently. The correlation coefficient between age effects estimated in the whole cohort and in sites 1/2 are 0.58/0.80 for *ϑ*_0_, 0.56/0.80 for *ϑ*_2_, 0.25/0.90 for *ϑ*_4_, 0.26/0.89 for *ϑ*_6_, respectively. The correlation coefficient for sex effects estimated in the whole cohort and in sites 1/2 are 0.66/0.68 for *ϑ*_0_, 0.69/0.64 for *ϑ*_2_, 0.50/0.81 for *ϑ*_4_, 0.51/0.80 for *ϑ*_6_, respectively.

**Figure 4:**
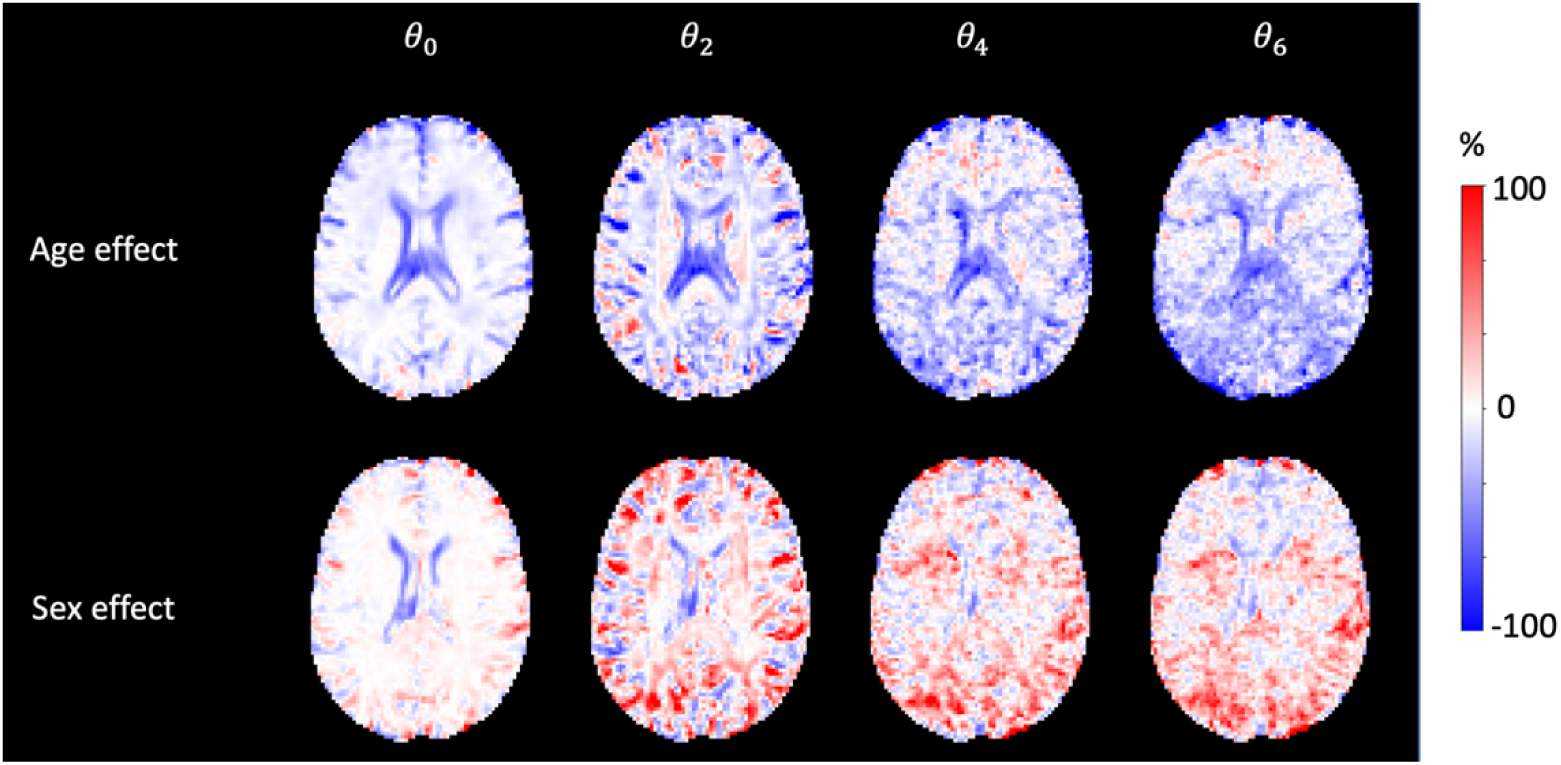
Percentage effect of age and sex estimated by RISH-GLM[Unmatched] in dataset 2. Age effects are prevalent at the interface between cerebrospinal fluid and white/grey matter for *ϑ*_0_, and in the white matter for *ϑ*_2_ and *ϑ*_4_. The age effect map for *ϑ*_6_ seems to be dominated by noise effects, given the low anatomical contrast, and the presence of clear stripes due to ghosting artefacts. Similar observations hold for sex effects.

In Figure 5, we evaluated the average FA values calculated in the white matter skeleton of the unmatched groups. Before harmonization, differences in average FA values equal to −4.8% (p≤0.05) between the two sites can be observed. Importantly, such differences can originate from both scanner-specific differences as well as from true biological effects given the large age span. Before harmonization, no correlation is observed between age and FA. In this situation, applying RISH[Unmatched] while ignoring age and sex effects reduces the average FA difference between the two groups to 0.1% (p=0.95), but suppresses the expected difference between two groups with largely different age distributions^35,36^. After harmonization with RISH-GLM[Unmatched], a relative difference in average FA values equal to 4.6% can be observed (p≤0.05), together with a linear negative relation between age and average FA that is in line with what was observed in Figure 2 (correlation coefficient rho = −0.64 vs −0.78 in Figure 2). Similar considerations hold also for MD, as shown in Supporting Information Fig. S2.

**Figure 5:**
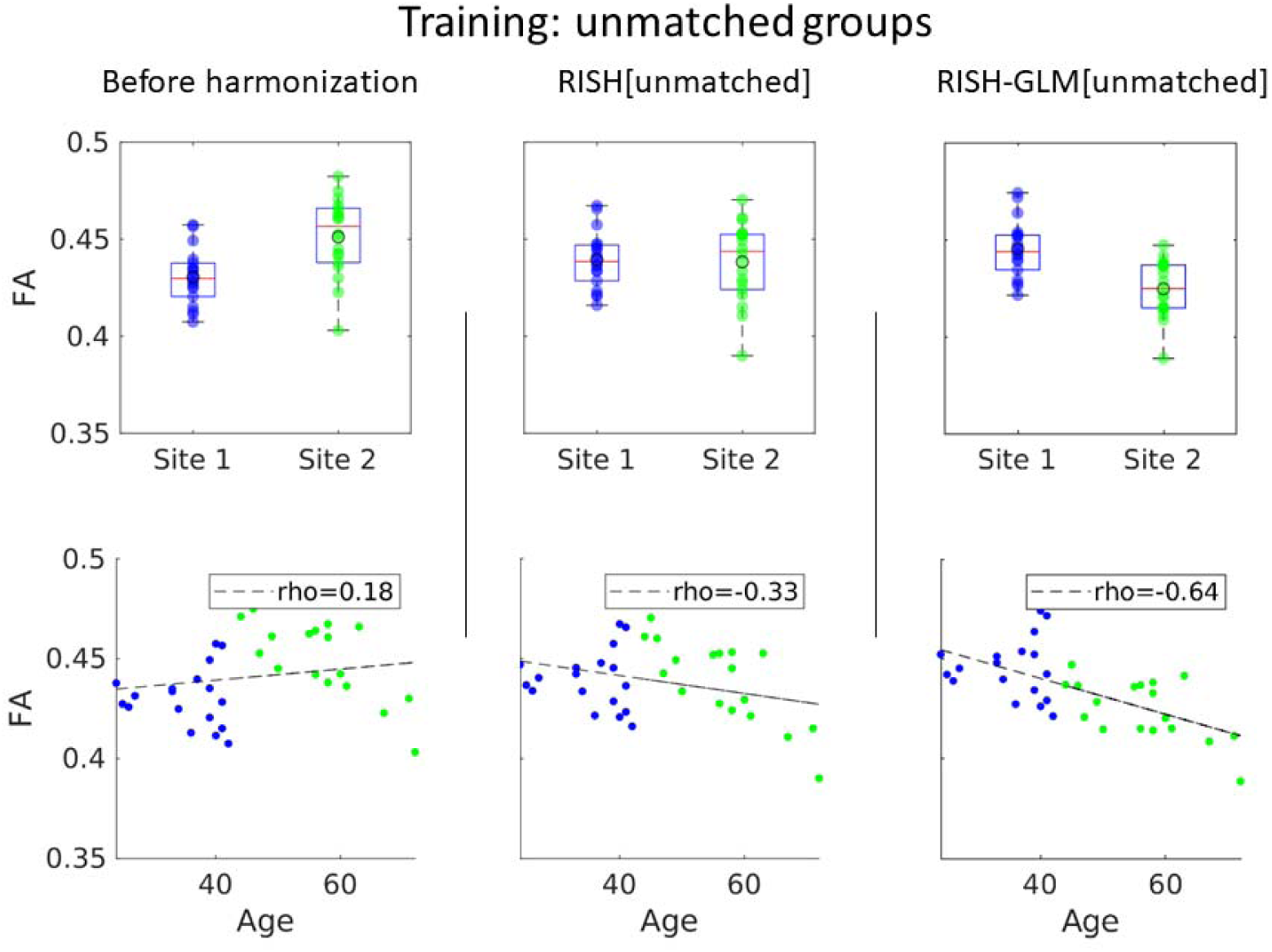
Boxplots of average FA values from two unmatched groups of healthy controls from Site1 and Site2 (dataset 2), before harmonization, and after harmonization with RISH and RISH-GLM. Before harmonization, an average difference in FA between the two groups can be observed. While this could be plausible given the unmatched nature of the two groups, there is an unexpected lack of correlation between age and FA before harmonization. Harmonization with RISH[Unmatched] removes the difference between the two groups. Consequently, a mild negative correlation between age and FA is observed, but the datapoints from both sites are not well distributed around the regression line, indicating a biased fit. After harmonization with RISH-GLM[Unmatched], a difference between the two groups in average FA values is observed, as expected, as well as a clear negative relation between age and FA.

Subsequently, we have applied RISH[Unmatched] and RISH-GLM[Unmatched] to harmonize the data of the 2 matched groups of Dataset 1, for which no average difference in FA values is expected after harmonization. The boxplots shown in Figure 6 showcase how RISH-GLM[Unmatched] can effectively remove the cross-site differences in FA between the two groups (average relative difference 0.5%, p=0.71), whereas RISH[Unmatched] does not (average relative difference −4.4%, p≤0.05). When looking at the relation between age and FA, the application of RISH-GLM[Unmatched] recovers the same age-FA relation observed also in Figure 2 and Figure 5. Similarly, RISH-GLM[Unmatched] effectively harmonizes also MD, as shown in Supporting Information Fig. S3, whereas RISH[Unmatched] does not.

**Figure 6:**
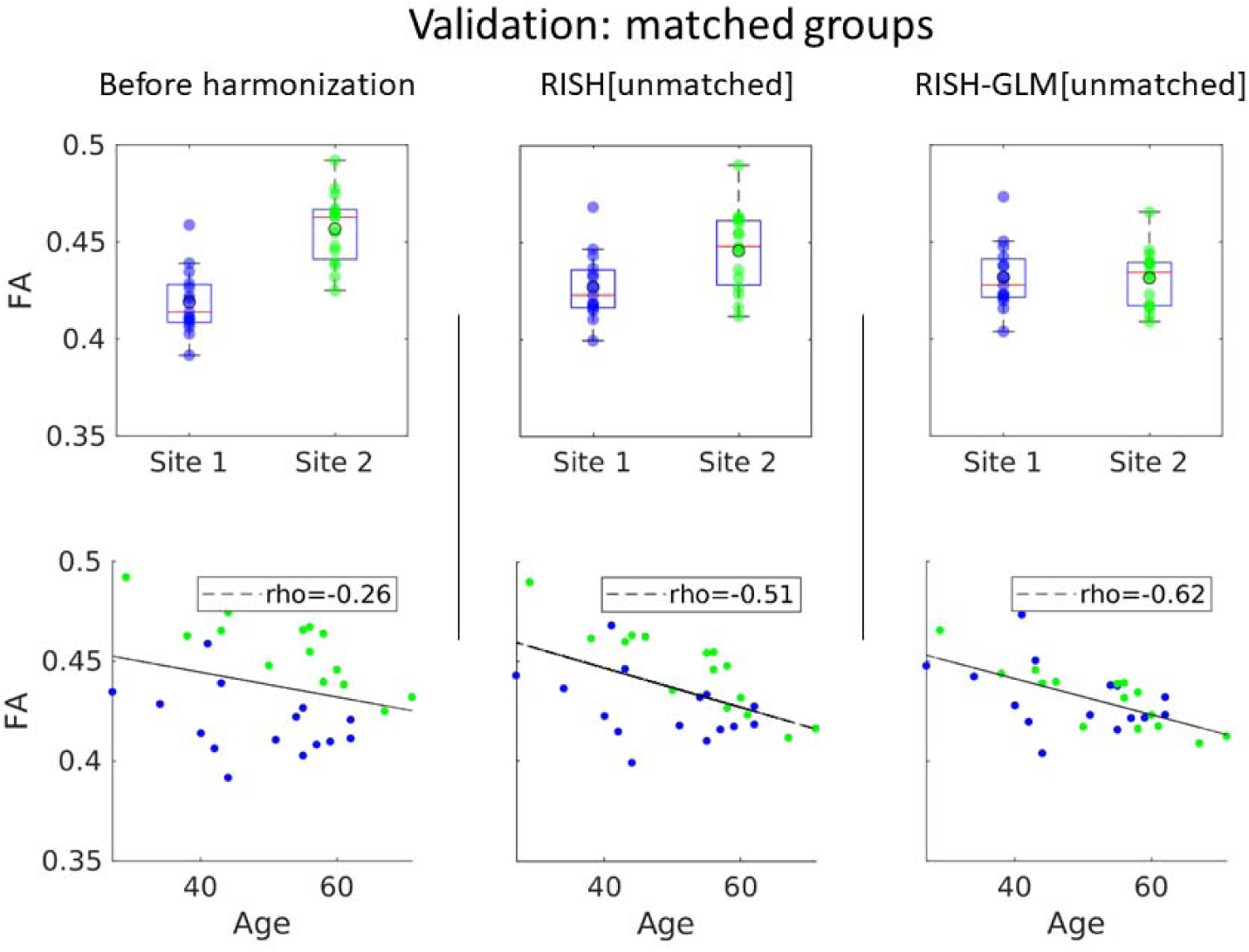
Boxplots of average FA values from two matched groups from Site1 and Site2 (dataset 2) before harmonization, and after applying RISH and RISH-GLM trained on unmatched data (Figure 5). No differences in average FA values are observed after harmonization with RISH-GLM, as expected for two matched groups. The application of RISH-GLM also allows to reveal the same negative correlation between age and FA observed in the previous figures.

### Experiment 3: One step harmonization

We investigated the ability of RISH-GLM to harmonize data from 3 sites in a single step. Example axial slices of the training scales derived with RISH and RISH-GLM between corresponding pairs of sites are shown in Supporting Information Fig. S4. Scaling maps computed from both methods have similar appearance, it can be appreciated how the scaling computed with RISH-GLM are higher on average than those computed with RISH. The presence of ghosting artefacts can be appreciated in the scaling maps *ϑ*_4_ and *ϑ*_6_, particularly with RISH-GLM. For this method, systematic artefacts can be learnt as part of cross-site differences, and can propagate to age and sex effect maps, as shown in Supporting Information Fig. S5. We subsequently evaluated differences in voxel-wise FA values between pairs of sites by means of permutations tests corrected by age and sex, obtaining the maps shown in Figure 7. Before harmonization, 32.1%, 0.25% and 29.6% of the brain voxels were statistically different when comparing Site 1 to Site 2, Site 1 to Site 3, and Site 2 to Site 3, respectively. After harmonization, the amount of significantly different voxels was reduced to 0.1%, 0.1% and 0.4%, respectively. Similar observations apply to MD, for which significant differences were observed in respectively 25.6%, 0%, and 31.6% of brain voxels when comparing sites in the same pairwise as previously reported. After harmonization, significant differences were reduced to 0.1%, 0%, 1.2% of brain voxels, respectively.

**Figure 7:**
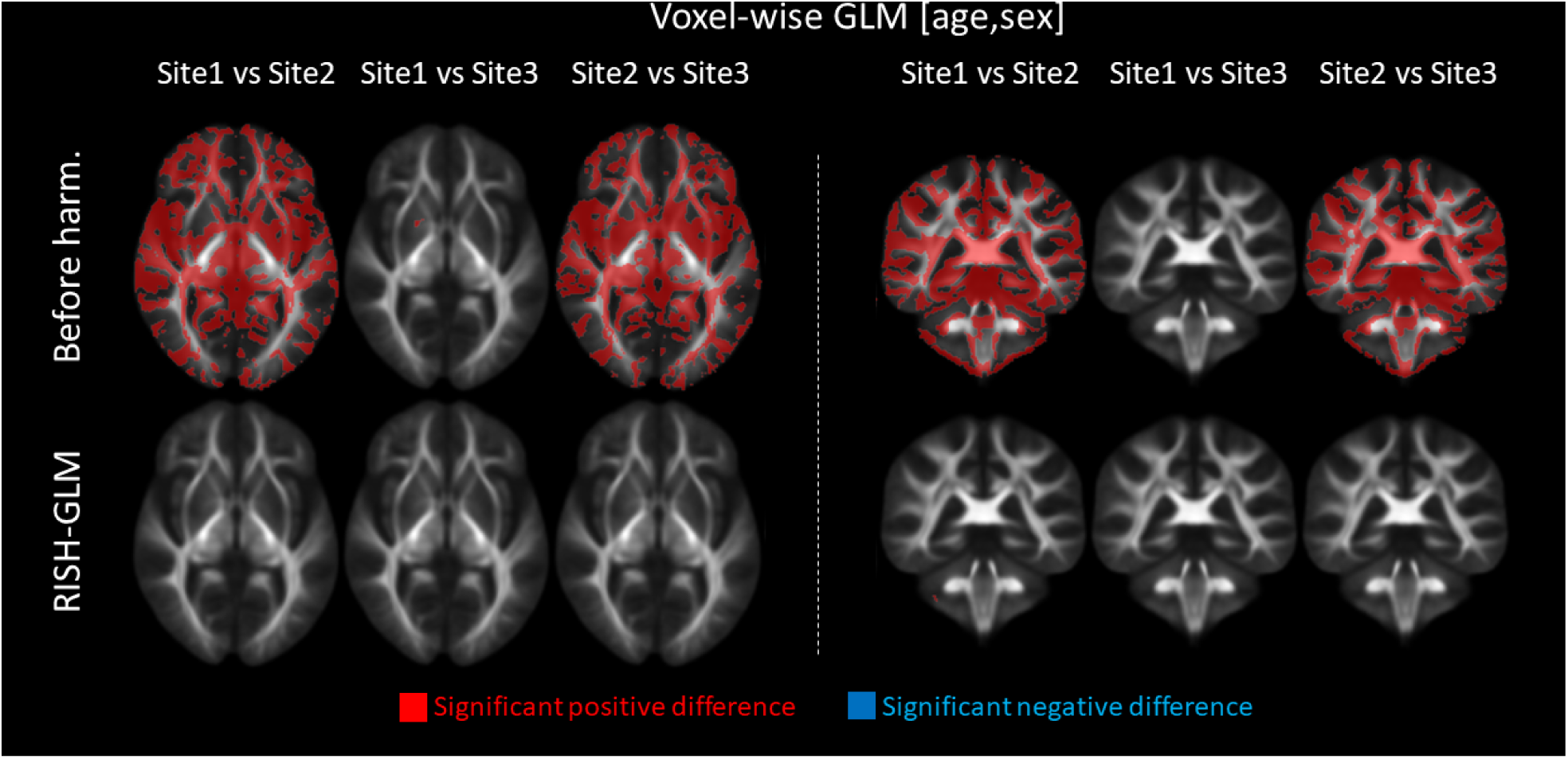
Example axial (left) and coronal (right) slices showing differences in average FA values at the voxel-level between pairs of sites of dataset 3 by applying a permutation test corrected for age and sex with a general linear model to three unmatched groups. Before harmonization, minimal to no differences are observed between Site1 and Site3. Conversely, extensive differences are observed between Site2 and the other sites. After harmonization with RISH-GLM in a single-step, most differences are effectively removed.

To validate the method, we have applied RISH-GLM to harmonize the independent dataset 4. Results, which are reported in Figure 8, demonstrate that RISH-GLM can effectively reduce the differences between all cohorts, and recover the same relation between age and FA/MD previously observed in Figure 2, Figure 5 and Supporting Information Fig. S6.

**Figure 8:**
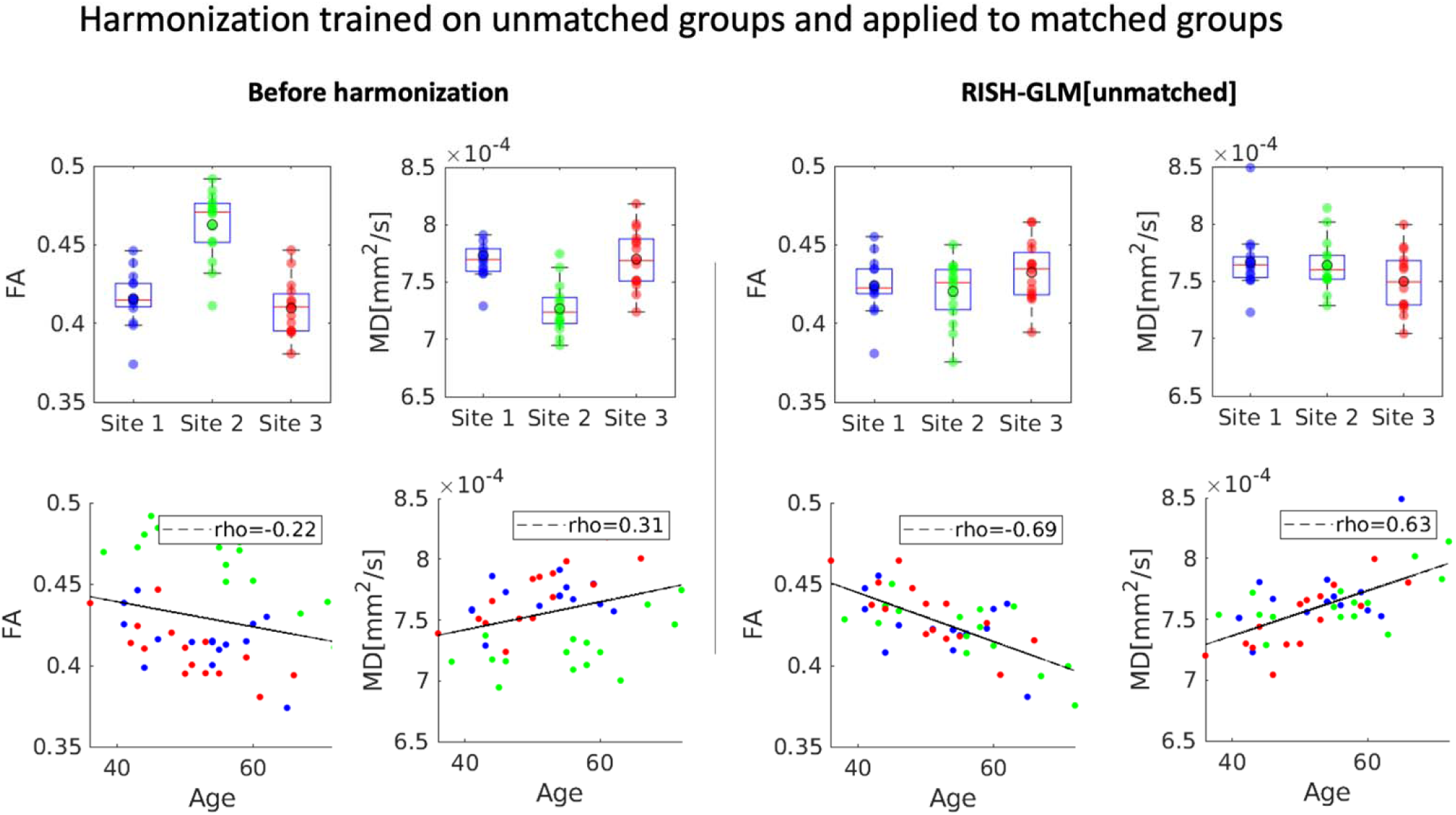
A comparison of average FA and MD values calculated in the white matter skeleton for three matched groups of subjects from Site1, Site2 and Site3 (dataset 4) before and after harmonization in a single step with RISH-GLM. After harmonization, no statistically significant difference between the three sites can be observed in both FA and MD, and values are well distributed around the regression line in the correlation plot between age and FA for all sites.

## Discussion

In this work, we have introduced a novel approach to perform RISH-based harmonization of the dMRI signal without need for training subjects matched at the group-level. Our approach does so by learning the voxel-wise effects of covariates on RISH scaling coefficients with a GLM (RISH-GLM). Our results demonstrate that RISH-GLM is effective in learning cross-site harmonization from unmatched training groups and can be used to harmonize data from multiple sites effectively in a single step.

The advent of dMRI harmonization methods such as RISH allows studies to overcome the barriers of single-site analyses, unharnessing the potential of retrospectively collected data to boost sample sizes. However, the applicability of such frameworks can be challenging when considering multiple cohorts selected with different inclusion criteria, because their requirement of matched training. Importantly, the results shown in Figure 5 and Figure 6 demonstrate that conventional RISH is unsuitable to learn cross-site harmonization with unmatched groups, as that would lead to a removal of both cross-site and biological effects (e.g., aging, Figure 5), or to the introduction of biases (Figure 6). RISH-GLM can effectively tackle this limitation by lifting the need for matched training groups, as demonstrated in Figure 6 and Figure 8.

The ability to learn cross-site harmonization without matched cohorts could support several efforts in both clinical and neuroscientific research. Firstly, RISH-GLM could simplify the harmonization of cohorts with only partly overlapping covariates of interest. This could be for example age, in the case of studies pooling retrospective cohorts with distinct age ranges to cover the whole lifespan. Secondly, it could support further disentangling scanner effects from confounding biological effects than conventional RISH, particularly when learning harmonization from groups which are matched on average but feature relevant variance in covariates. In the study of dementia, for example, population cohorts, memory clinic cohorts, and population with different injury aetiologias could be potentially harmonized while accounting for key factors that could otherwise bias the harmonization process, such as differences in education, ethnicity, sex distributions, exposure to specific risk factors, lesion burden, etc.

While retrospective data is arguably the main application goal of post-processing dMRI harmonization methods such as RISH-GLM, they could prove advantageous also to support prospective studies. In general, accurate matching of acquisition hardware and sequence parameters can already reduce cross-site dMRI variability, as shown by previous initiatives in frontotemporal^37^ and vascular dementia^38^. Yet, subtle differences between vendors, calibration, and characteristics of individual scanners might still dictate the existence of cross-site differences. Furthermore, even when prospective matching is achieved successfully at the start of a study, subsequent hardware changes and software updates might still introduce acquisition effects in long multiyear prospective studies. In the future, software versions and coil updates could be considered as covariates of interest in RISH-GLM to attempt limiting their effect on the acquired data.

When comparing scaling maps computed with RISH-GLM using matched datasets to those computed with RISH (Figure 1), local relative differences up to 20% can be observed close to the WM-GM and WM-CSF interfaces. This is likely due to taking into account age and sex effects during estimation, and suggests that such features may be locally relevant even in matched datasets. This observation is supported by the fact that relative differences between unmatched groups, shown in Fig. 3, exhibit spatial patterns that are consistent with those of Fig. 1, but with larger amplitude. Despite such local differences, when comparing FA and MD of matched groups (Fig. 2 and Supporting Information Fig. S2), we did not observe any relevant group-difference with both RISH and RISH-GLM. However, our analysis clearly focused on the WM skeleton, whereas most differences between RISH and RISH-GLM are located close to WM/GM interface. As such, future studies should investigate the potential of RISH and RISH- GLM on more sophisticated analyses such as fiber tractography, where the transition of fibers between tissue types is crucial to achieve meaningful results.

It is important to acknowledge some limitations of this study. As explained in the methods section, RISH- GLM is based on assumptions that might not always hold true. An important assumption is the existence of a linear relation between covariates of interest and RISH features, at least within the range of covariates used for training. In the case of age, for example, little is known about its exact relation to RISH features. Furthermore, RISH-GLM has primarily been designed to account for systematic cross-site biases, and its ability to capture non-systematic effects leading to different subject ranking across sites^39^ remains unclear. Evaluating such aspects and validating the assumptions of RISH-GLM throughout the lifespan will require dedicated studies, or an additional validation in cohorts including so-called travelling-heads, i.e., groups of subjects travelling across sites and scanned with highly controlled experimental conditions to ensure minimal variance of their measurements across sites^13,39^ (e.g., ideally even considering confounders such as time of the day^40^). Nevertheless, we observe that RISH-GLM is effective at recovering consistent relations between age and FA throughout the experiments of this work. Of note, the assumption of linearity between covariates and RISH features does not imply the need for linearity between covariates and derived metrics such as FA, given that RISH features are non-linear descriptors of the diffusion signal. This is particularly relevant as previous studies^35,41^ of diffusion tensor imaging metrics across the lifespan have demonstrated the existence of a quadratic – locally linear – relation between age and fractional anisotropy, for example. Another assumption of RISH-GLM is that the effects of covariates is similar across training cohorts. As a consequence, training directly with non-homogeneous groups – such as patients with different pathologies – is discouraged. In this study, we used data from a retrospective cohort of individuals that were initially marked as at risk for frontotemporal dementia, which might have introduced a selection bias towards effects not explicitly accounted for. Furthermore, we only focused on modelling the effect of age and sex, and did not consider other relevant confounders such as education.

In conclusion, we have introduced a novel framework to learn cross-site harmonization while accounting for covariates of interests, and demonstrated its effectiveness to learn harmonization between two age unmatched groups with an average age difference of over two decades.

## Acknowledgements

This study is funded by the Bluefield Project to cure FTD. The research of ADL is also supported by independent funding from Alzheimer Nederland (WE-03-2022-11).

## Code availability statement

The code used to perform RISH and RISH-GLM harmonization is openly available at https://github.com/delucaal/RISH-GLM.

---

Figure S1: Percentage effect of age and sex estimated by RISH-GLM[Matched]. Age effects are prevalent at the interface between cerebrospinal fluid and white/grey matter for *ϑ*_0_, and in the white matter for *ϑ*_2_ and *ϑ*_4_. The age effect map for *ϑ*_6_ seems to be dominated by noise effects, given the low anatomical contrast, and the presence of clear stripes due to ghosting artefacts. Similar observations hold for sex effects.

Figure S2: The top row shows boxplots of average MD values in the white matter skeleton per site, before harmonization (first column), after harmonization with RISH (middle column) and with RISH-GLM (last column). The second row shows scatterplots of the same average MD values as a function of age. Harmonization with both RISH and RISH-GLM was trained with group level-matched subjects.

Figure S3: Boxplots of average MD values from two unmatched groups of healthy controls from Site1 and Site2, before harmonization, and after harmonization with RISH and RISH-GLM. Before harmonization an unexpected negative relation between age and MD is observed. Harmonization with RISH removes any relation between age and MD. After harmonization with RISH-GLM[Unmatched], a positive relation between age and MD is observed, as expected based on previous literature.

Figure S4: Scaling maps calculated between pairs of sites with RISH, and in one single step with RISH-GLM on all three sites considered in Experiment 3.

Figure S5: Percentage effect of age and sex on RISH features of different orders as determined with RISH-GLM.

Figure S6: Boxplots of average MD values from two matched groups from Site1 and Site2 before harmonization, and after applying RISH and RISH-GLM trained on unmatched data (Figure 5). No differences in average MD values are observed after harmonization with RISH- GLM, as expected for two matched groups. The application of RISH-GLM also allows to reveal the same positive correlation between age and MD observed in the previous figures.

## Notes

### Competing Interest Statement

The authors have declared no competing interest.

### Summary of Updates

The manuscript has been revised taking into account the comments received from reviewers after submission to MRM. Methods and discussion have been clarified, and results now demonstrate that RISH-GLM also harmonizes MD. The code of the framework has been made available and linked in the text.

