## Supporting Information for "Cross-site harmonization of diffusion MRI data without matched training subjects"

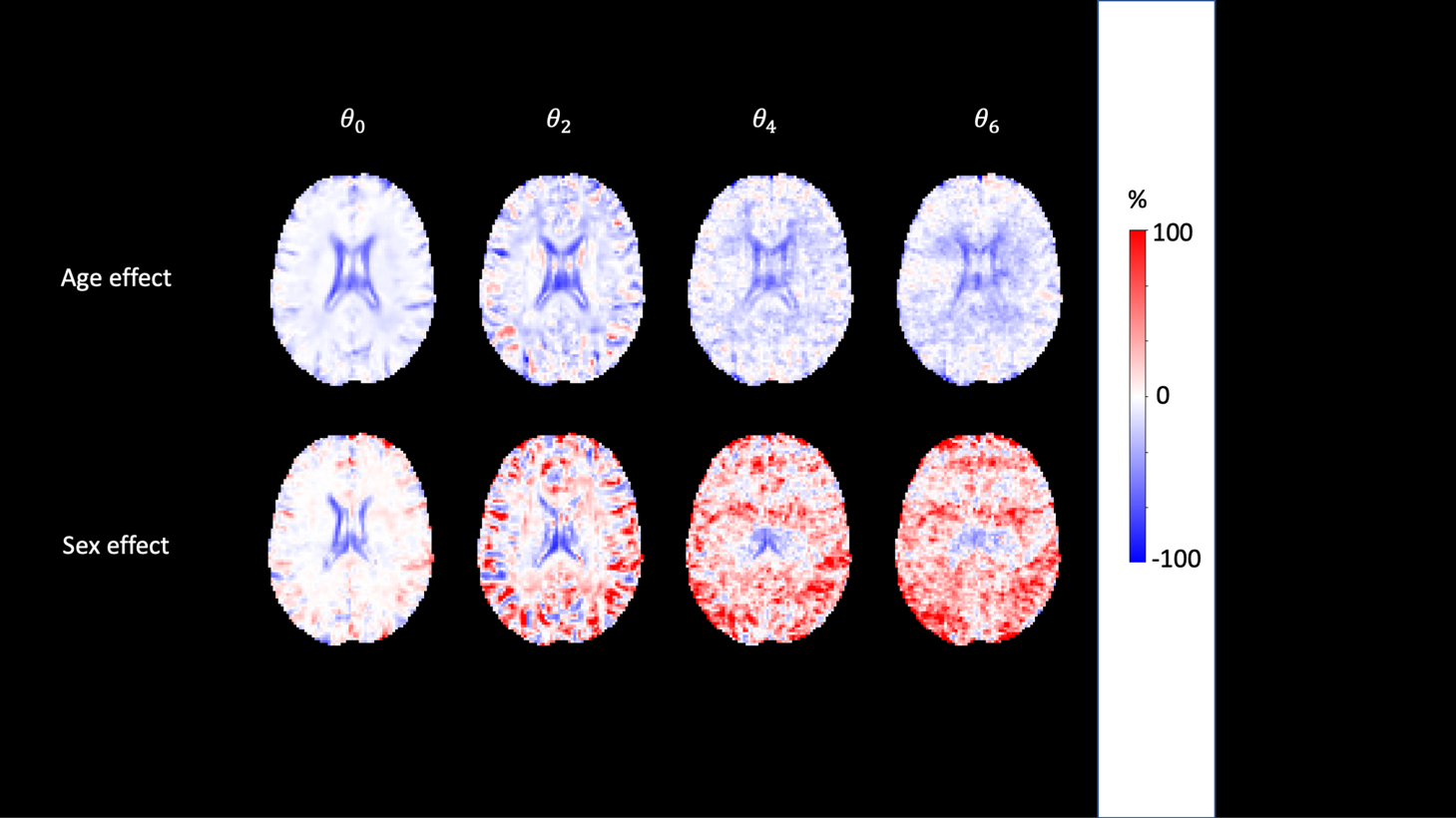

Figure S1: Percentage effect of age and sex estimated by RISH-GLM[Matched]. Age effects are prevalent at the interface between cerebrospinal fluid and white/grey matter for $\vartheta_{0}$, and in the white matter for $\vartheta_{2}$ and $\vartheta_{4}$. The age effect map for $\vartheta_{6}$ seems to be dominated by noise effects, given the low anatomical contrast, and the presence of clear stripes due to ghosting artefacts. Similar observations hold for sex effects.

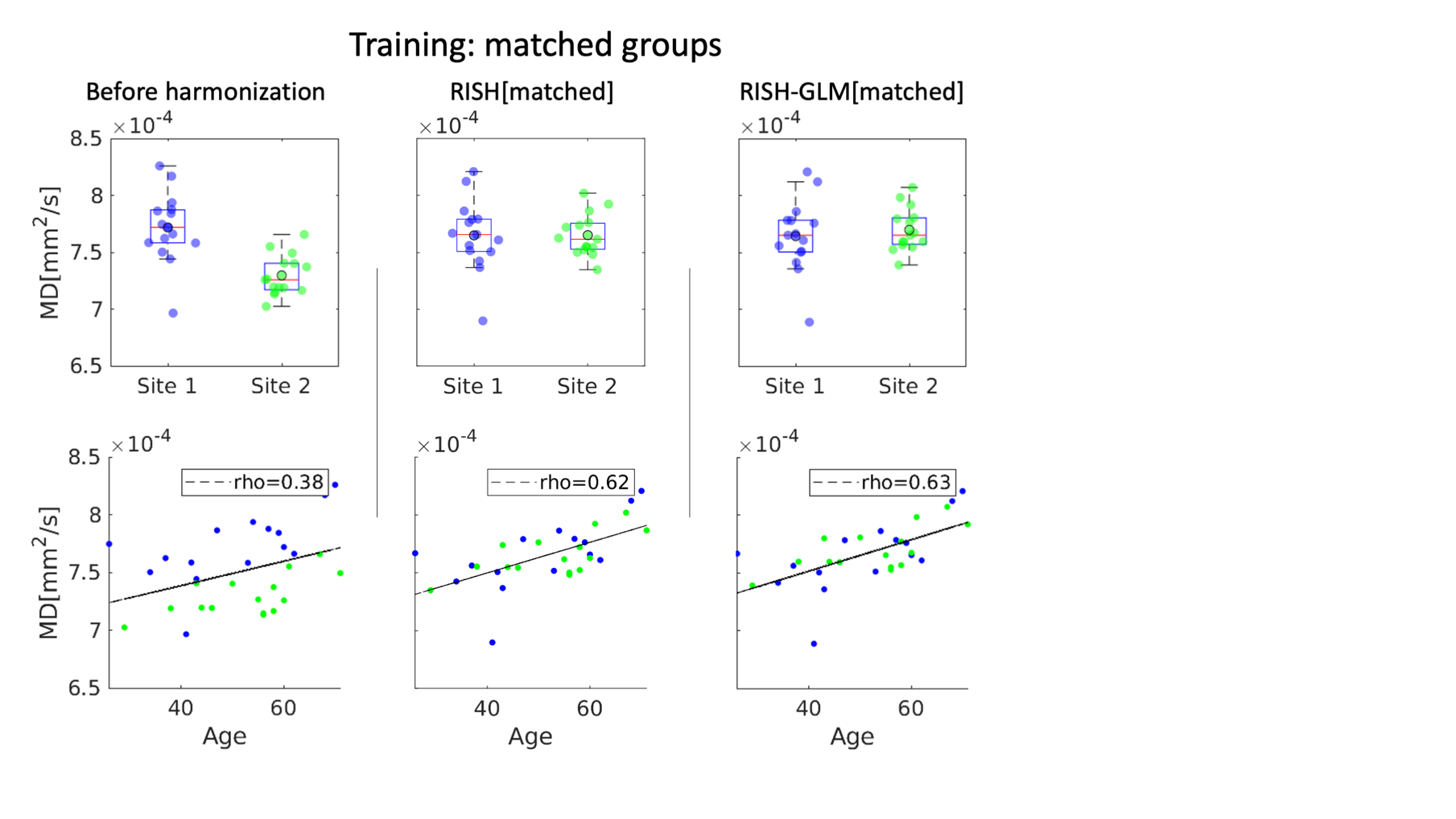

Figure S2: The top row shows boxplots of average MD values in the white matter skeleton per site, before harmonization (first column), after harmonization with RISH (middle column) and with RISH-GLM (last column). The second row shows scatterplots of the same average MD values as a function of age. Harmonization with both RISH and RISH-GLM was trained with group level-matched subjects.

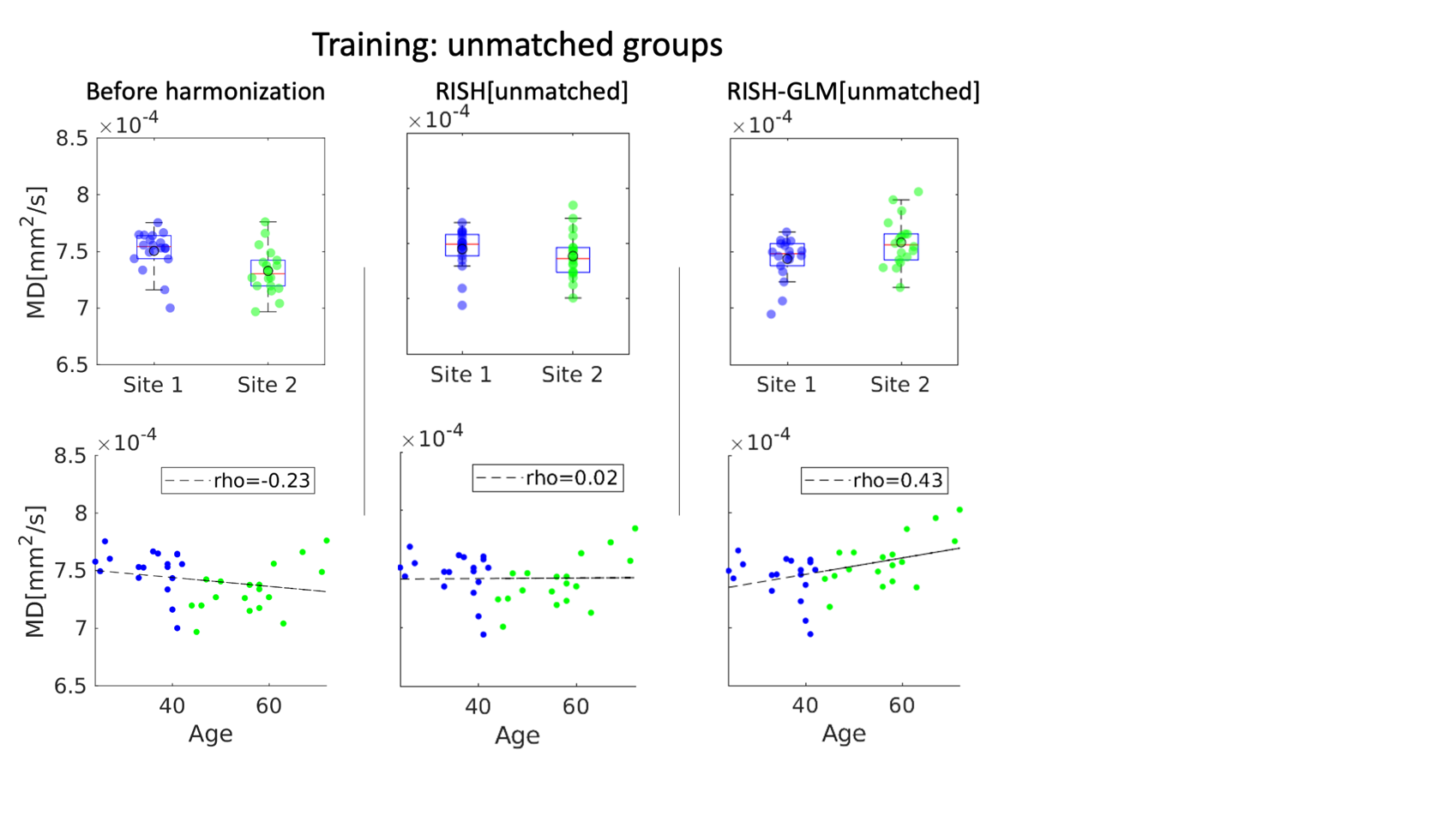

Figure S3: Boxplots of average MD values from two unmatched groups of healthy controls from Site1 and Site2, before harmonization, and after harmonization with RISH and RISH-GLM. Before harmonization an unexpected negative relation between age and MD is observed. Harmonization with RISH removes any relation between age and MD. After harmonization with RISH-GLM[Unmatched], a positive relation between age and MD is observed, as expected based on previous literature.

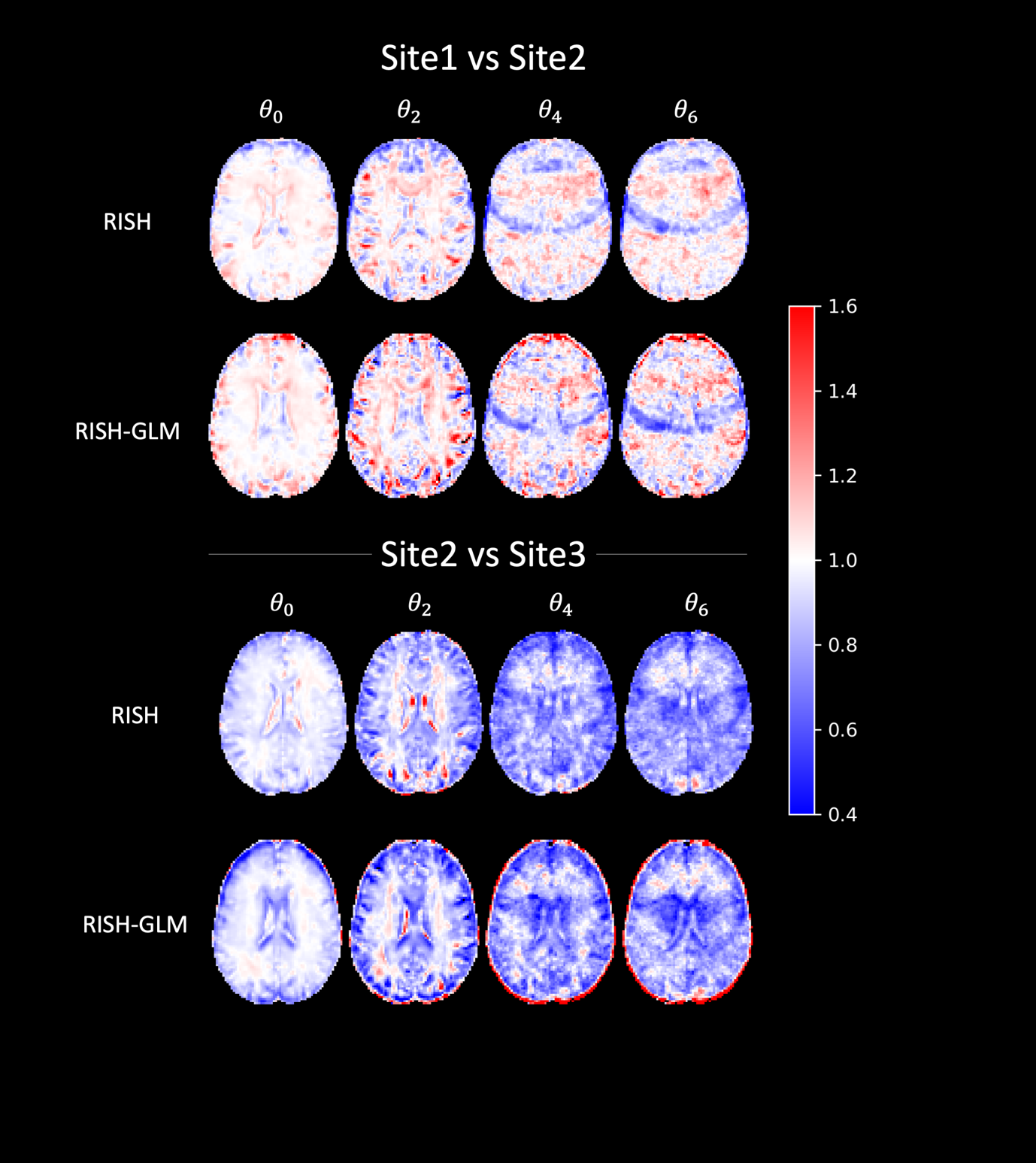

Figure S4: Scaling maps calculated between pairs of sites with RISH, and in one single step with RISH-GLM on all three sites considered in Experiment 3.

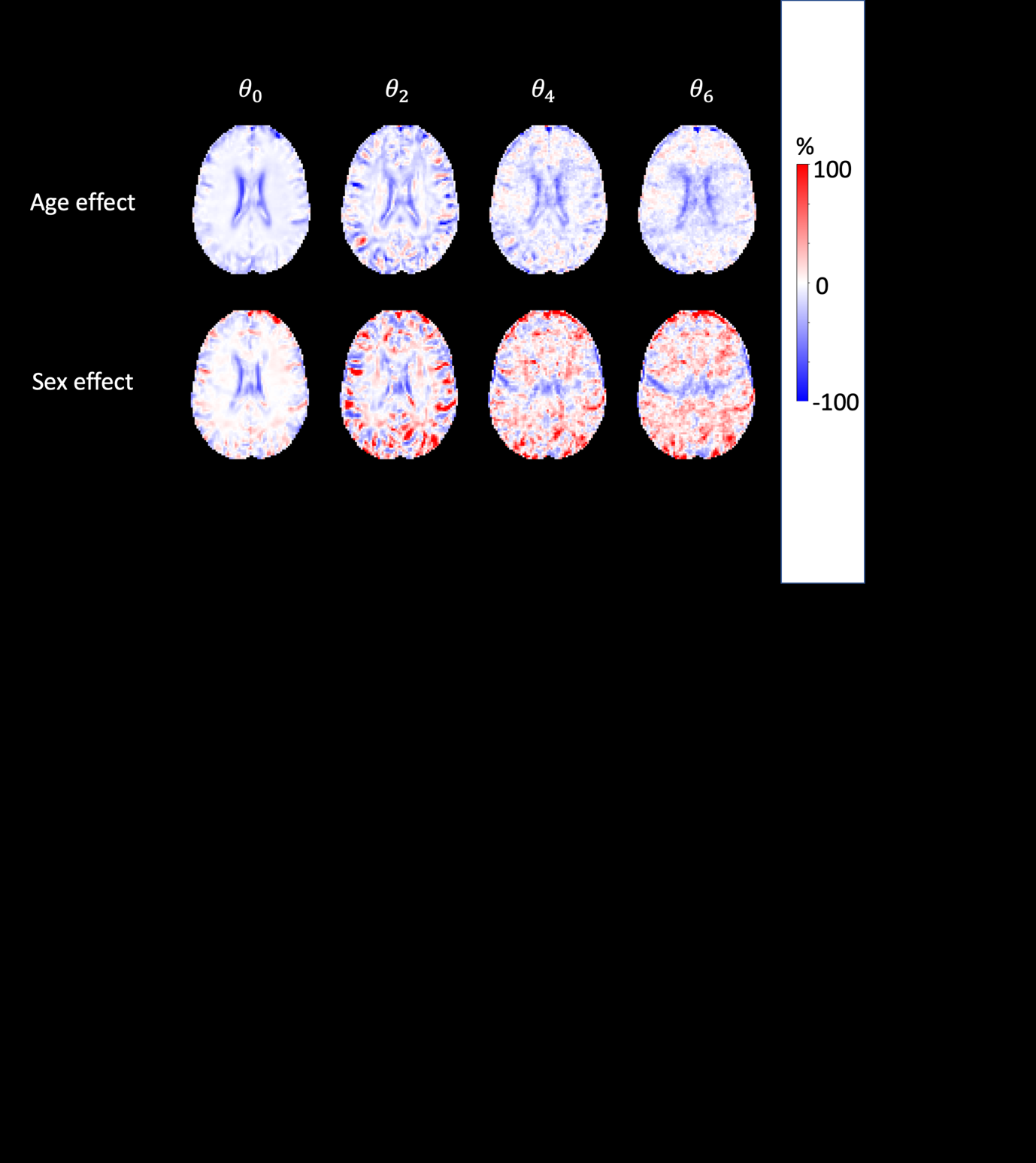

Figure S5: Percentage effect of age and sex on RISH features of different orders as determined with RISH-GLM.

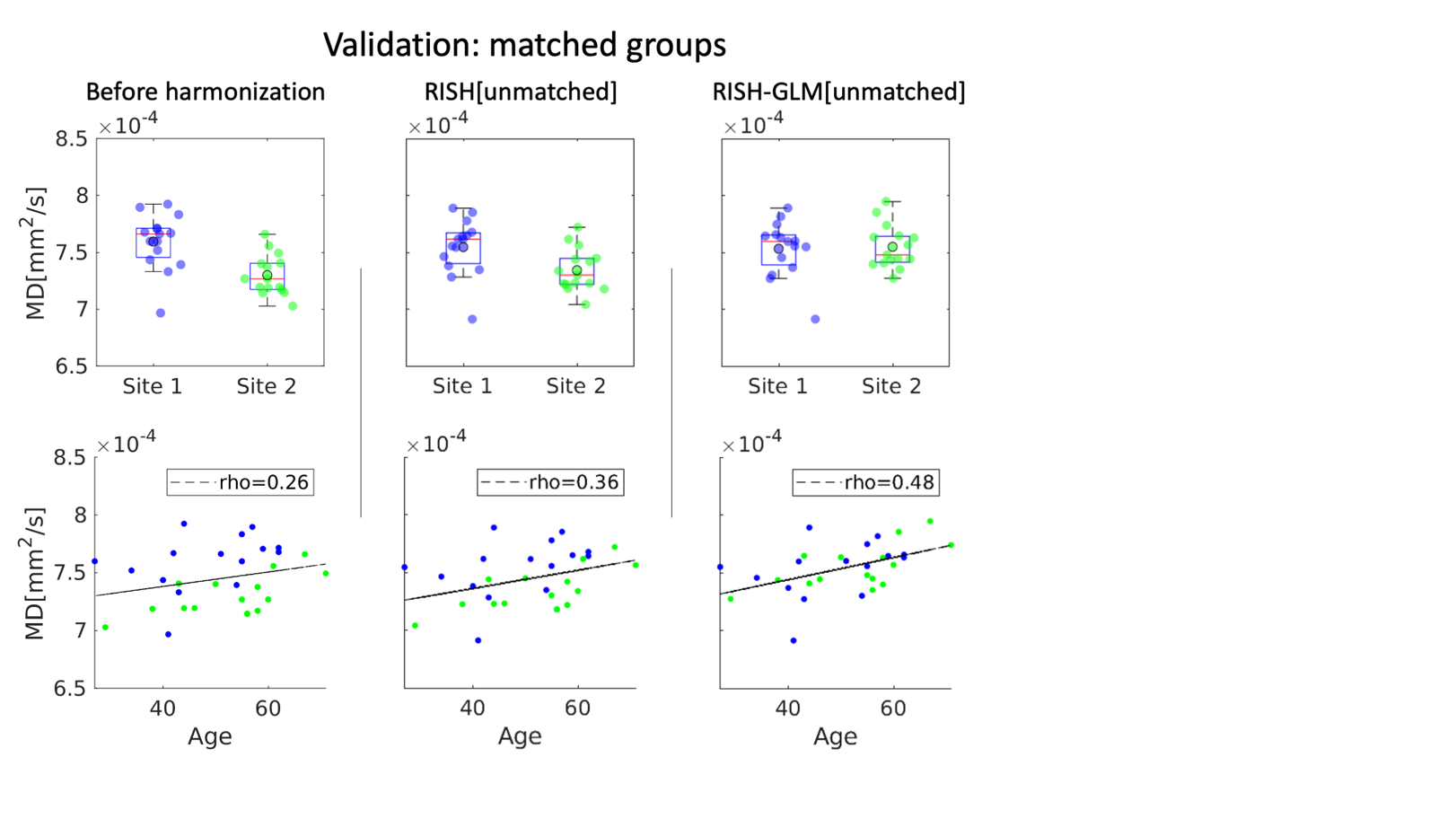

Figure S6: Boxplots of average MD values from two matched groups from Site1 and Site2 before harmonization, and after applying RISH and RISH-GLM trained on unmatched data (Figure 5). No differences in average MD values are observed after harmonization with RISH-GLM, as expected for two matched groups. The application of RISH-GLM also allows to reveal the same positive correlation between age and MD observed in the previous figures.
